# Easy Two-Photon Image-Scanning Microscopy With Spad Array And Blind Image Reconstruction

**DOI:** 10.1101/563288

**Authors:** S. V. Koho, E. Slenders, G. Tortarolo, M. Castello, M. Buttafava, F. Villa, E. Tcarenkova, M. Ameloot, P. Bianchini, C.J.R. Sheppard, A. Diaspro, A. Tosi, G. Vicidomini

## Abstract

Two-photon excitation (2PE) laser scanning microscopy is the imaging modality of choice when one desires to work with thick biological samples. However, its spatial resolution is poor, below confocal laser scanning microscopy. Here, we propose a straightforward implementation of 2PE image scanning microscopy (2PE-ISM) that, by leveraging our recently introduced ISM platform – based on a new single-photon avalanche diode (SPAD) array detector – coupled with a novel blind image reconstruction method, is shown to improve the effective resolution, as well as the overall image quality of 2PE microscopy. Indeed, in stark contrast to conventional single-point detectors, SPAD array detectors give access to the images of any excited scanning region, from which it is possible to decode information about the aberrations/distortions – occurring during imaging – able to substantially improve the reconstruction. Most importantly, our 2PE-ISM implementation requires no calibration or other input from the user; it works like any familiar two-photon system, but produces higher resolution images deep into thick samples. In our view, this novel implementation is the key for making 2PE-ISM mainstream.

---

With the explosion of super-resolution methods [1, 2], fluorescence microscopy research has been experiencing somewhat of a renaissance. A plethora of new optical microscopy techniques have been introduced, some able to “move” the spatial resolution limit beyond the diffraction barrier [3, 4], and others – usually referred to as nanoscopy or diffraction-unlimited techniques – able to break such a barrier and reach a resolution of only few nanometers [5, 6, 7]. However, despite the big technological advances and promising proof-of-concept demonstrations, nanoscopy techniques have not been able to replace conventional microscopy and traditional super-resolution methods, such as confocal laser scanning microscopy (CLSM) [8] and structured-illumination microscopy (SIM) [9] – as go-to imaging tools in pre-clinical research. These techniques are reliable, simple, familiar, highly compatible with all kinds of fluorescence labels and work well with many types of samples – whereas current nanoscopes fall short on at least some of these characteristics.

SIM encompasses a collection of super-resolution implementations that make use of structured excitation light [9]. In contrast to nanoscopy, SIM does not require any special sample preparation [10]. In its original form [3], SIM was implemented in a wide-field microscope, by producing – *via* interference – a striped illumination pattern onto the sample with a line spacing close to the diffraction limit, thereby shifting high-frequency information of the sample into the pass-band of the optical system. Because the excitation pattern is diffraction limited, the maximum resolution gain obtainable with linear SIM techniques is – from a cut-off frequency point-of-view – a factor of two with respect to conventional microscopy [3].

Similarly to SIM, CLSM maintains the large sample-compatibility, and the two-fold resolution gain can be obtained by completely closing the pinhole [11]. However, such an enhancement is only theoretical, since it comes at the cost of extremely low signal-to-noise ratio (SNR). This problem can be overcome by image-scanning microscopy (ISM) [12]. In a nutshell, starting from a conventional CLSM architecture, the pinhole is removed and the single-point detector is substituted by a detector array, which collect an “image” of each excited region, i.e., for each scanning position. Because each element of the detector array acts as a pinhole, but no photons reaching the image-plane are rejected, by fusing all the scanned images (one image *per* detector’s element) – *via* deconvolution or pixel-reassignment (PR) – a super-resolved and high-SNR image is reconstructed.

Image-scanning microscopy was theoretically proposed in the 80s [13, 4], but an effective implementation – from here traditional ISM – was only recently achieved thanks to the development of fast detector arrays (bandwidth ≫ kHz) – faster than the typical CLSM pixel-dwell time, such as the Airyscan [14] and our tailor-made single-photon-avalanche-diode (SPAD) array module [15]. However, to compensate for the lack of fast detector arrays, early multiple-spots [16, 17] or all-optical/re-scanned [18, 19] ISM implementations – based on a conventional camera – have been introduced. In all-optical ISM, the image reconstruction is achieved by optically enlarging the final image by a fixed factor – usually two – with respect to the laser scanning grid. Later, multiple-spots and all-optical ISM implementations have been combined together [20, 21]. All these implementations are sometimes classified as SIM techniques, since they use structured excitation/illumination (spots instead of stripes), wide-field architectures and conventional cameras. Henceforth, traditional ISM is sometimes called spot-scanning SIM [22]. Roughly speaking, SIM and ISM can be considered the “two faces of the same coin”: the former uses structured-illumination, the second structured-detection, but both achieve similar resolution enhancement. However, because of the complexity in generating patterned illumination deep into a sample, traditional SIM imaging is usually constrained to a few tens of micrometers in depth [2], also when combined with adaptive optics approaches [23]. On the other hand, thanks to the point-scanning architecture, the traditional ISM is more compatible with thick samples.

Initially all ISM implementations have been implemented with one-photon excitation (1PE), which limits the practically attainable imaging depth in thick biological samples, due to extensive scattering of the illumination light. More recently, to address this issue, both all-optical and traditional ISM implementations employing two-photon excitation (2PE) have been proposed [24, 25, 26, 27, 28]. 2PE is well known to allow a much deeper penetration, and the infrared excitation light is also less phototoxic, as biological samples do not commonly absorb it [29, 30, 31]. Moreover, the two-photon excitation process improves the optical sectioning capability to the extent that a separate pinhole is not usually necessary [32].

However, the current all-optical 2PE-ISM implementations – as for the 1PE counterpart – require significant changes to the regular point-scanning microscopy architecture and are not robust against sub-optimal or variable imaging conditions. On the other hand, the traditional ISM implementations give access to important information about the imaging conditions/distortions, as recently demonstrated by us [15]. In a nutshell, traditional ISM effectively provides access to the images of the single excited regions, whilst all-optical ISM averages-out this information because the camera integrated during the scanning - as in regular laser-scanning microscopy where the single-point detector averages light across its whole sensitive region. As a matter of fact, this novel “spatial” information can be leveraged to reconstruct super-resolved and high-SNR images, even when working under sub-optimal and variable imaging conditions, such as in deep imaging. This advantage, to the best of our knowledge, has not bee previously explored in the context of 2PE-ISM. An approach to compensate for imaging distortions in multiple-spots ISM has been proposed [33], but it requires tedious pre-calibration of the system – prior to every imaging session. Furthermore, such calibration is error-prone, as the imaging conditions in the actual sample, especially in depth, cannot be expected to be similar to the calibration sample.

Here, inspired by our recent SPAD array-based ISM platform [15], we present a calibration-free, robust and straightforward 2PE-ISM implementation, which can be implemented on any regular point-scanning microscope. In fact, as shown in (Fig. S. 1), our implementation is essentially just a simple upgrade to a confocal laser scanning microscope, with only a couple of additional lenses and our new SPAD array detector. Similarly to our one-photon excitation ISM implementation [15], we automatically record all the raw scanned images – without speed concerns and large data overhead that hampered the early multiple-spots ISM implementations. We propose two fully automatic and adaptive image reconstruction methods, aimed to decode the imaging conditions/distortions and to generate the super-resolved and high-SNR image without any calibration measurements or prior knowledge of the sample or microscope configuration. While the first improves the robustness of our previous adaptive-PR method, the second combines the benefits of the adaptive-PR and the blind deconvolution reconstruction approaches. We show how our algorithms allow improving the resolution – typically to the theoretical limit – in a variety of samples.

In our view, the simple optical configuration and the novel image reconstruction methods are rather important steps towards a wider adoption of ISM, as they entirely hide its complexity; ideally the end user would just see a regular confocal or 2PE microscope, with improved performances.

## Results

### 2PE-ISM *via* Pixel-Reassignment

In traditional image-scanning microscopy, i.e., implemented with a fast detector array, the most straightforward method to fuse the scanned images – one for each detector element – is pixel-reassignment (PR) (Fig. 1(a)). The method is based on the simple observation that any element of the detector array provides – in good approximation – a confocal image which is a copy of the other scanned images, but shifted in space. In particular, each scanned image is an “ideal” confocal image: each detector element acts as a pinhole, which effective size – the physical size of the detector element back-projected on the image plane – is ≪ 1 Airy unit. Thereby, successfully shifting-back all the scanned images – with respect to a reference – before summing them up results in a high-SNR image with an optical resolution, i.e. the full-width at half-maximum (FWHM) of the associated point-spread-function, equal to the “ideal” confocal counterpart. We first performed simulations [34] to quantify the resolution enhancement achieved by implementing 2PE-ISM with our 5 *×* 5 -element SPAD array detector and with the PR approach (Fig. 1(b-d)). Fundamental aspects of the PR approach are the shift-vectors used to register all the scanned images. According to the PR theory [35] and using the image of the central element as reference, the shift-vector **s**_*i*_ = (*x, y*) associated with the detector element *i* is based on simple geometrical considerations: (i) the shift-vector direction equals the line joining the centers of the considered element and the central one; (ii) the module is proportional to the PR factor *α*_*PR*_, the magnification of the system *M*, and the physical distance between the element and the central one *d*_*i*_, namely 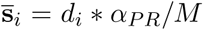 (iii) assuming identical excitation and emission wavelength, the pixel reassignment factor is equal to 0.5 – notably, this condition is satisfied only in the case of 1PE and no Stokes-shift; (iv) in the case of non-negligible Stokes-shift and 2PE a theoretical PR factor can be estimated [36]. However, in our simulation, we obtained the shift-vectors *s*_*i*_ by co-aligning the peak intensity points of all element’s PSFs. As expected, our approach generates shift-vectors similar to the theoretical method – described above – for elements close (*≤* 1 AU) to the central reference (PR factors reported in Fig. S. 2).

**Fig. 1.**
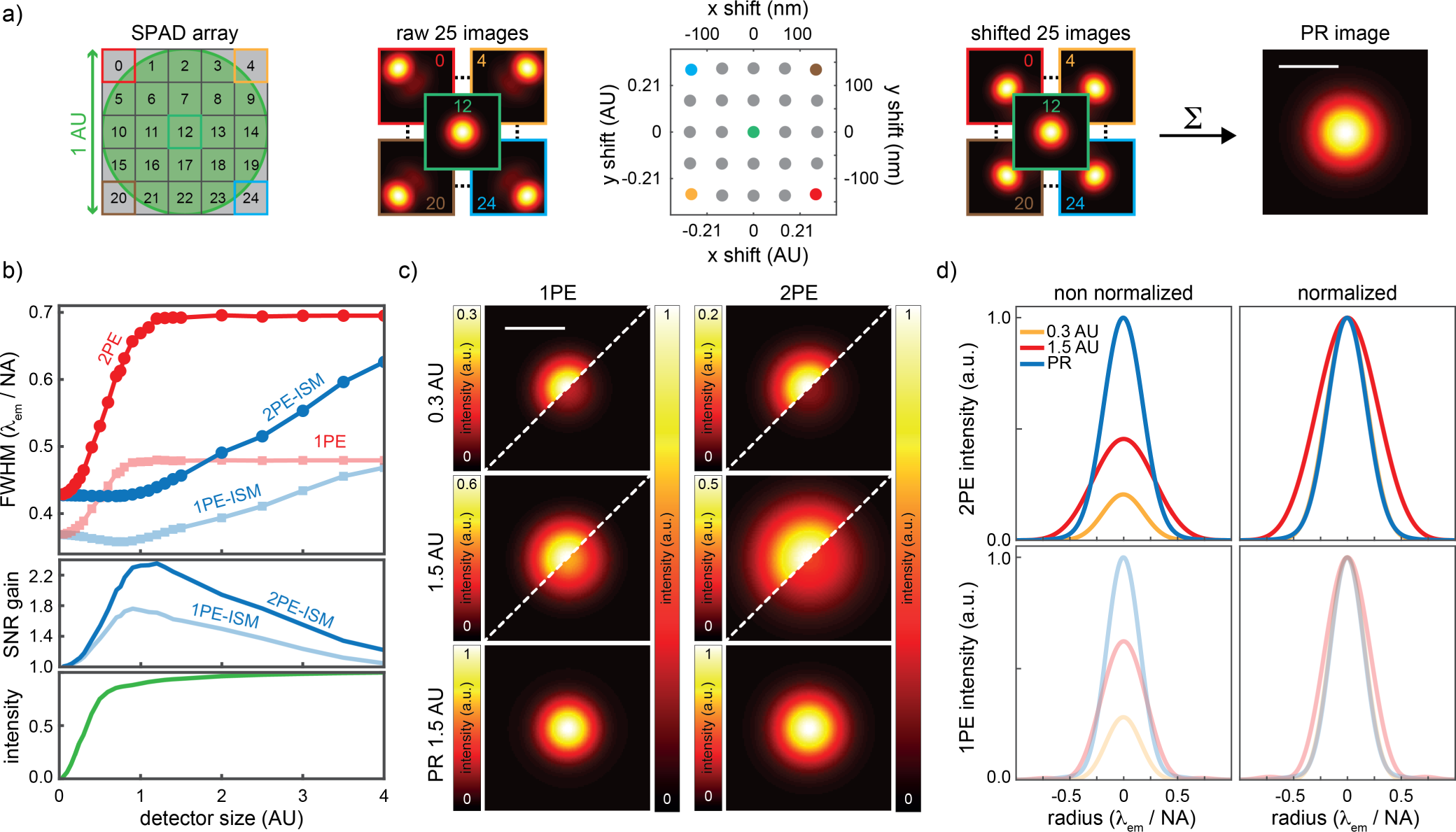
Point-Spread-Function simulation for 2PE-ISM. (a) Schematic representation of the simulation to calculate the PSFs of ISM based on the pixel-reassignment method. The effective detector size depends on the magnification of the microscope system and is usually expressed in Airy Unit, i.e., 1 AU = 0.61*×λ*_*em*_*/NA*, where *λ*_*em*_ is the wavelength of the fluorescence light and NA is the objective numerical aperture. In this simulation we used: NA = 1.4, *λ*_*em*_ = 525 nm. Each element of the detector array acts as a shifted pinhole, thus 25 PSFs have been calculated using a rigorous model for the focused emission/excitation light intensity distribution (*λ*_*exc-*2*PE*_ = 950 nm, *λ*_*exc-*1*PE*_ = 488 nm, oil objective lens). The shift-vectors for the simulation are calculated by assuming that an optimal shift should co-aligned all PSFs at their maximum intensity. Finally, all the registered PSFs are summed to produce the PSF of ISM. (b, top) Optical resolution (i.e, FWHM of the PSF) comparison between confocal (i.e.,the PSFs are summed without any shift) and ISM as a function of the detector size, for both the 1PE and 2PE case. (b, middle). Gain in SNR (i.e., the ratio between the peak intensities of the ISM and the confocal PSFs) as a function of the detector size, for both the 1PE and 2PE case. (b, bottom) Fraction of fluorescence recorded – with respect to the total emission – as a function of the detector size; as expected, there is no difference between 1PE and 2PE, neither confocal or ISM. (c) Side-by-side comparison of the PSFs for the different imaging configurations. Here, 1 AU is assumed as detector size. Both intensity -normalised and -unnormalised versions are reported. (d) Radial intensity profiles for the PSF shown in c). Scale bars: 1 *µm*

Since the number of elements of our SPAD array is limited, an optimal magnification *M* of the system (i.e., an optimal effective detector size) needs to be found in order to collect the majority of photons focused on the image plane and, at the same time, maintain an effective structured detection. Whith this in mind, we calculated the PSF for both 2PE-ISM and 2PE as a function of the effective SPAD array size (in Airy unit of the focused fluorescence light). Figure 1(b, top, bottom) shows the optical resolution (FWHM of the PSF) and the total fluorescence intensity, respectively. Large detector sizes (*>* 1.5 A.U.) destroy the resolution enhancement expected from ISM, and small detector sizes (*<* 1 A.U.) lose a substantial part of fluorescence signal. The range 1-1.5 A.U. grants a resolution enhancement of 1.6 and a negligible 10% loss of the fluorescence signal. Notably, the improvement obtained by ISM in 2PE is stronger compared to the one-photon excitation counterpart 1(c,d): (i) the resolution gain for the 2PE case is *∼* 1.6 versus *∼* 1.4 for one-photon excitation; (ii) the super-brightness effect (i.e., the peak intensity increase between the ISM and the pinhole-less PSFs) is *∼* 2.2 for 2PE and *∼* 1.8 for one-photon excitation (Fig. 1(b, middle)).

### 2PE-ISM *via* Adaptive Pixel-Reassignment and Image Reconstruction

The PR reconstruction is very sensitive to the shift-vectors. Above we described the theoretical approach largely used to derive the shift-vectors, which is also at the base of the all-optical ISM implementations. However, similar to the approach used in our simulation, it does not consider realistic elements such as optical aberrations and system misalignments. For these reasons, an automatic (i.e., model-free and calibration-free) method able to estimate realistic shift-vectors is highly desirable, especially for those experiments in which the imaging conditions change significantly and continuously, e.g., 3D imaging in thick samples. To satisfy this need, we have recently introduced a method that we called adaptive-PR, which is able to calculate the optimal shift-vectors for the image reconstruction directly from the scanned images. Here, we improve the robustness of this method and we show its synergistic integration with blind image reconstruction/deconvolution. Adaptive-PR consists of two tasks: image registration and image fusion. Image registration is needed to derive the shift-vectors, aligning the 25 scanned images/sub-images (Fig. 2b) into a common coordinate system; image fusion is used to combine the registered sub-images, summing them up or *via* more complex operations, into a single result. Notably, if the registration step is bypassed, the sum of all the scanned images results in the conventional pinhole-less 2PE image (Fig. 2a). In our previous work [15], we estimated the shift-vectors by means of a phase-correlation approach implemented into the Fourier domain. Here, we propose a new iterative image registration method, based on software originally developed for three-dimensional tomographic STED microscopy [37]. In image registration one tries to find a spatial transformation that aligns the details in two images as closely as possible. In iterative image registration, as illustrated in (Fig. 2c) the search for the optimal spatial transformation is considered an optimisation problem: one image, called *fixed image*, is used as a reference, while a second image, called *moving image*, is translated until the details in the two images match. The images are considered to match when a chosen similarity *metric* reaches its maximum value. In our ISM registration implementation, the central detector image is always used as the *fixed image* and the 24 other images are sequentially registered with it. After the shift-vectors for all the 24 scanned images have been calculated, the images are shifted and then added together to produce the adaptive-PR ISM (APR-ISM) image. However, both in ISM and SIM some form of deconvolution or frequency domain filtering (re-weighting) is commonly used in order to maximise the effective resolution (e.g. [24, 34, 25, 3, 20]). To this end, as illustrated in (Fig. 2d)), we apply our blind Fourier ring correlation (FRC) based Wiener filtering approach [38] to the APR-ISM image; no prior knowledge of the point-spread-function (PSF) is needed, but it is estimated directly from the APR-ISM result image. In essence, the FRC analysis is applied on the APR-ISM image to extract the resolution, i.e., the cut-off frequency of the image, from which we derive the FWHM for a Gaussian PSF. Henceforth, we denote the combination of APR-ISM with blind deconvolution/Wiener filtering as APR-ISM^*b*+^. The image shifting and Wiener filtering could be alternatively combined in a single step, as is done in SIM [3, 39] and as we proposed in quadrant ISM [34]. In essence, similar to multi-image Richardson-Lucy deconvolution [15], the shift-vectors are directly encoded in the PSFs used in the reconstruction algorithm. Here, we keep the two tasks separated, because the FRC approach that we used to estimate the PSF is sensitive to noise: the estimation of the PSF for the APR-ISM image is more robust than the estimation of the PSFs for each single scanned image.

**Fig. 2.**
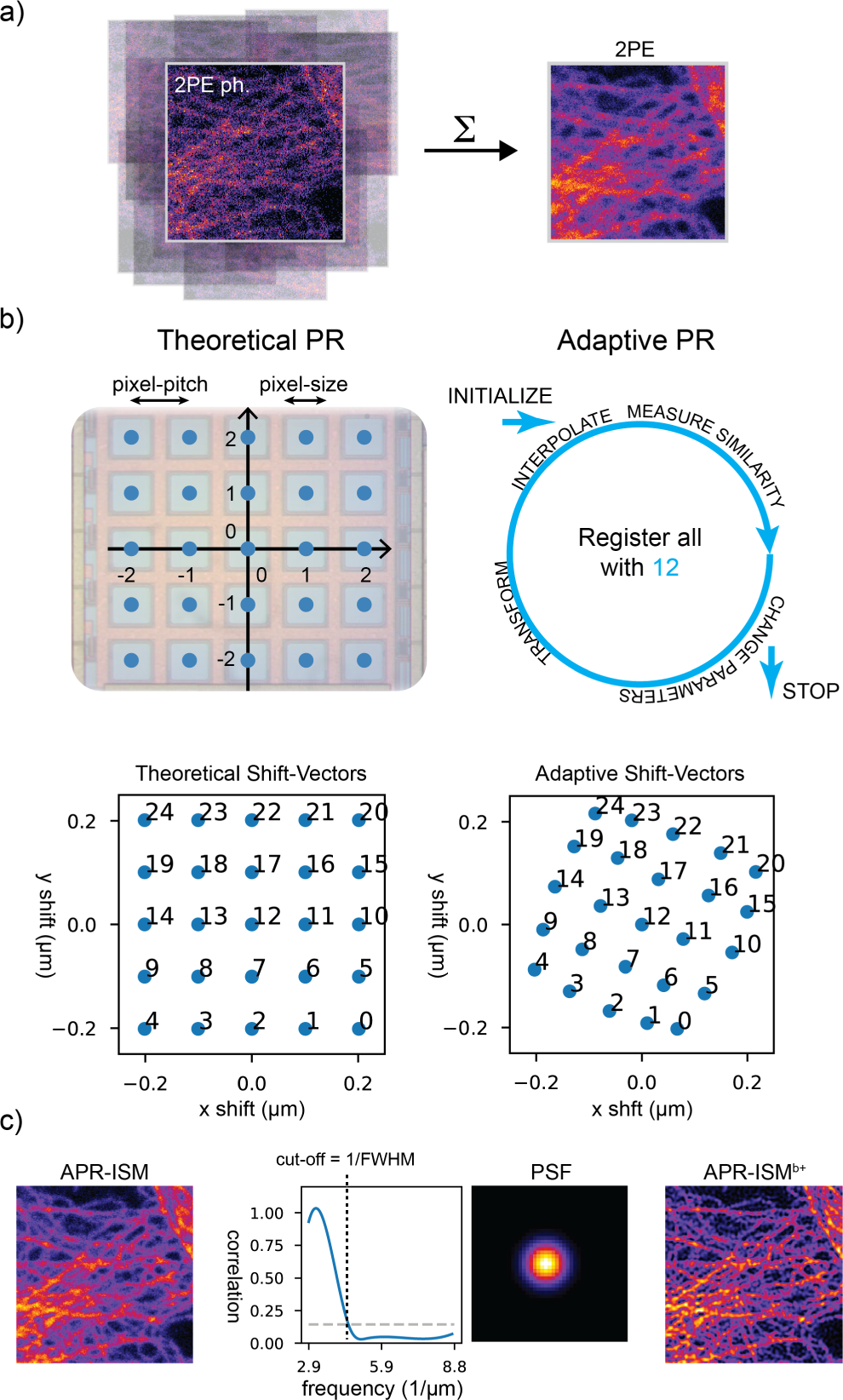
ISM image reconstruction methods at a glace. (a) At the end of a scan, an image, shifted with respect to all the others, is associated with each pixel 0-24 of the SPAD array (Fig. 1). By summing the 25 images, the regular 2PE image is obtained. (b) For pixel-reassignment ISM image reconstruction one first needs to determine the image shifts. The theoretical shifts matrix **s** is obtained by scaling the 5 *×* 5 meshgrid **gr** representing the SPAD array with the pixel pitch *p*_*p*_ = 75 *µm*, the magnification of the microscope *M* = 500, and the pixel-reassignment factor *α*_*PR*_, equal to 0.5 in this case, i.e., **s** = **gr** *\* p*_*p*_ *\* α*_*PR*_*/M*. For the adaptive PR, iterative image registration method is applied to align all the individual images with the image from the central detector element (12). The PR-ISM reconstruction is obtained by shifting and summing the 25 scanned images. (c) for the (APR-)ISM^*b*+^ a blind Wiener filter is applied to the reassignment result; the PSF is estimated from the ISM reassignment result image with FRC.

In addition to the APR-ISM^*b*+^ reconstruction method, we also propose a simple algorithm that allows constructing the ISM result image point-by-point during data-acquisition, similarly to regular confocal or 2PE microscopes (Note S 1). This is possible because our SPAD array detector has no frame rate, and thus every single photon can in real time be assigned to its correct spatial location. However, in this case one needs to know the shift-vectors from the beginning of the scanning, for example from previous acquisitions. Further, no sub-pixel shifts can be used, but a re-sampling of the reconstructed image can be used.

### Benchmark measurements using test samples

In order to get an idea of the performance of our 2PE-ISM system, we took some benchmark measurements under *quasi* aberration-free imaging conditions (at the coverslip interface), with two very common microscope test samples: i) a sample of 100 *nm* yellow-green carboxylate modified fluorescent nanospheres, and ii) a sample of fixed HeLa cells stained with an alpha–tubulin antibody and with a Star488 secondary antibody.

In (Fig. 3 a) we compare four images of the nanoparticle sample: the regular two-photon image (2PE), the confocal two-photon image (2PE ph.), the ISM image after our adaptive-PR (APR-ISM) method and the Wiener filtered APR-ISM image (APR-ISM^*b*+^). The 2PE image was obtained by summing the signal from all SPAD array elements, ignoring the registration step, whereas the 2PE ph. image corresponds to the image registered by the central element (pixel no 12 in Fig. 2 b); the pixel itself works as a pinhole of approximately 0.5 Airy units in size. For the APR-ISM image, the 25 sub-images were first registered and then summed. The ISM shift vectors obtained with image registration are shown in a (x,y) scatter plot in (Fig. 3 a). Interestingly, while the shift-vectors form a grid-like structure, even with a basic sample like beads the grid is somewhat tilted, probably due to aberrations/misalignment. The pixel no. 24 in the SPAD array used for the 2PE experiments is more noisy than the others, which appears to compromise the image registration with the sparse beads sample. As shown with the FRC analysis (Fig. 3 a), the APR-ISM method improves the resolution by a factor of *∼* 1.7, which is further improved to a factor of *∼* 3 with our blind Wiener filter, applied on the APR-ISM image (APR-ISM^*b*+^). The numerical resolution values obtained with FRC measurements on the nanoparticle sample (2PE: 295 nm, APR-ISM: 179 nm and APR-ISM^*b*+^: 79 nm) correspond quite well with the theoretical FWHM values (FWHM_2*PE*_ = 263 nm and FWHM_2*PEph*_ = FWHM_2*PE-ISM*_ = 147 nm from our PSF simulation (see Fig. 1)) Although the regular 2PE appears to perform sub-optimally, which can be explained by the low SNR, APR-ISM almost reaches the theoretical limit of the closed pinhole 2PE (2PE ph.) and 2PE-ISM.

Considering the fixed cell sample, as shown in (Fig. 3 b), the resolution of the APR-ISM image is improved by a factor of *∼* 1.6 with respect to the regular 2PE image counterpart; it is further enhanced by a factor of *∼* 2.5 thanks to Wiener filtering. However, it is worth noticing that the retrieved resolution values are different than the ones obtained with the nanoparticles sample (2PE: 373 nm, APR-ISM: 240 nm and APR-ISM^*b*+^: 110 nm); the 2PE resolution with the cell sample is close to the Abbe’s diffraction limit (*d*_*min*_ = *λ*_2*PE*_*/*(2NA) = 339 *nm*, for *λ*_2*PE*_ = 950 nm, NA = 1.4; quadratic dependency with the excitation intensity is not considered), and the APR-ISM resolution values are in good agreement with previous reports on non-linear ISM [24, 25]. Comparison between the adaptive PR method and the theoretical PR method (for different PR factors) is also reported (Fig. S 3), showing the superior result of the adaptive method, also in a *quasi* ideal imaging condition.

**Fig. 3.**
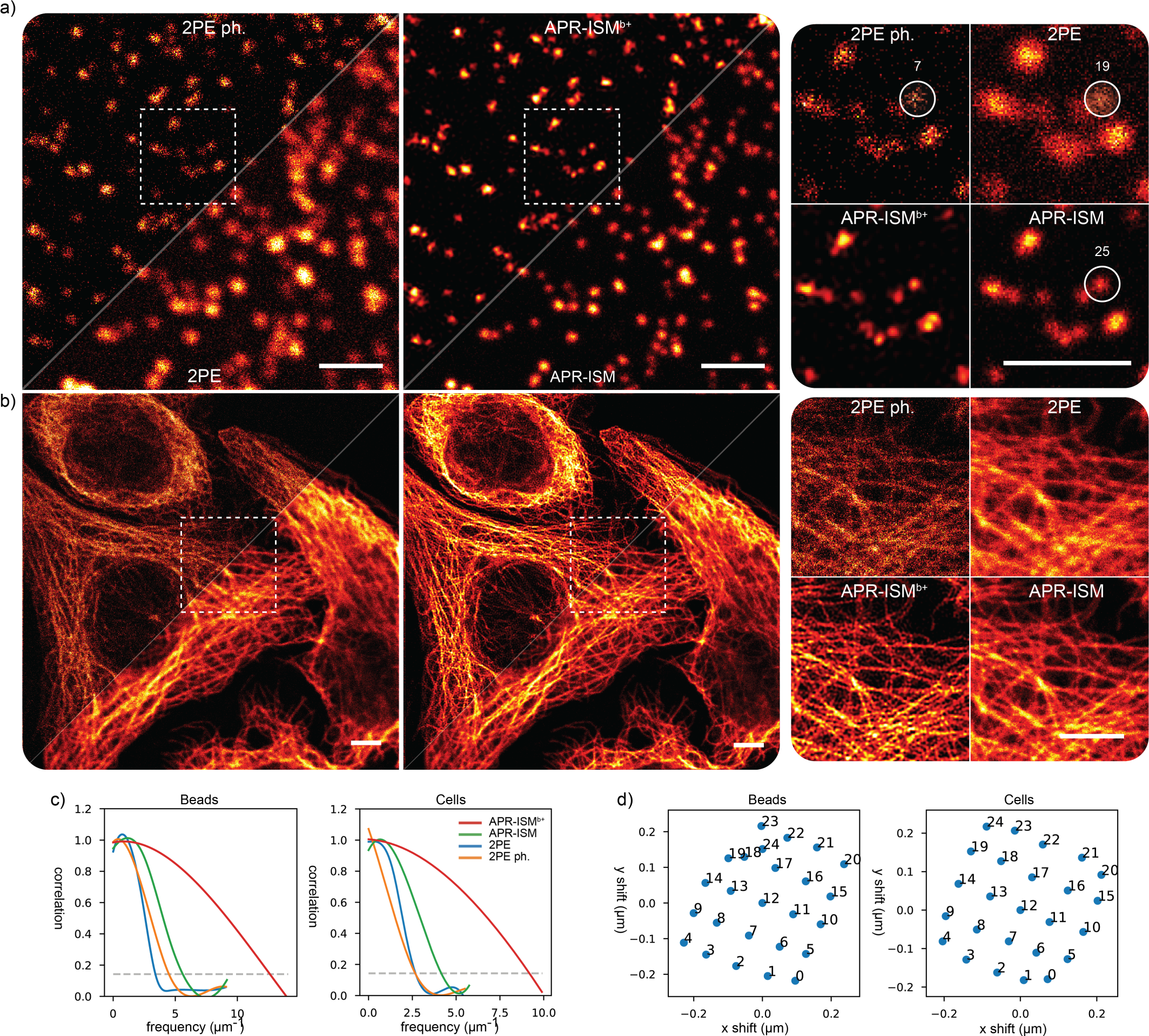
2PE-ISM imaging with simple test samples. (a,b) Side-by-side comparison of 2PE microscopy, confocal 2PE (2PE ph.) microscopy, PR-ISM, and deconvolved PR-ISM (PR-ISM^*b*+^) for imaging of 100 nm sized fluorescent nanoparticles (a) and HeLa cells (b). Insets show the magnified views of the regions inside the white dashed boxes. Peak intensities for the fluorescent nanoparticles highlighted in the white circle are reported. (c) The FRC measures show a nearly three-fold resolution improvement in APR-ISM^*b*+^ with respect to regular 2PE. (d) The shift-vectors are very similar for both of the experiments, and form a somewhat tilted grid. The shifts are a bit smaller for the cell image, suggesting a larger PSF size, which is consistent with the FRC resolution measurements. All images and insets are normalised to the respective maximum intensity values. Scale bars: 2 *µm* (a); 4 *µm* (b)

We then tried to push the resolution in APR-ISM further by iterative Richardson-Lucy (RL) deconvolution (Fig. S 4). Although the RL is able to considerably boost the contrast, no major gains are made in terms of quantitative resolution. We show that our real-time pixel reassignment algorithm produces good results with the HeLa cell image (Fig. S 5), with similar level of details that were observed in (Fig.3 b)). A strong signal-to-noise ratio (SNR) gain is observed with respect to the 2PE ph. image: this is expected considering the strongly adverse effect on the SNR in 2PE microscopy caused by the introduciton of an optical pinhole (2PE ph.). While the features clearly get smaller, as suggested by the theory (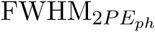 *versus* FWHM_2*PE*_), the effective resolution values are actually worse than in regular 2PE. This underlines the strength of ISM, and helps to understand why optical pinholes are not typically implemented in 2PE microscopes.

### Diving into a hazy brain slice

A multi-photon microscope arguably is most at home with thick samples that are hard or impossible to image with regular fluorescence microscopy methods, and thus we decided to test our 2PE-ISM in such samples. A partially optically-cleared mouse brain that expresses YFP in neurons was imaged at different penetration depths, up-to the maximum working distance of the microscope objective, *∼* 140 *µm*. The 2PE-ISM is able to maintain rich details and good contrast all the way up to the maximum working distance (Fig. 4 a). Whilst in regular 2PE the YFP appears distributed uniformly inside the neurons, 2PE-ISM is able to reveal a peripheral distribution, characteristic that is more and more evident in APR-ISM^*b*+^. Furthermore, adaptive PR-ISM^*b*+^ provides a constant image quality across all the imaging depths, by maintaining a spatial resolution of *∼* 140 *nm*, whereas the resolution in regular 2PE images degrades from 480 nm to 530 nm as a function of depth, as can be expected (see FRC measurements in Fig. 4 b); this corresponds to a resolution improvement of a factor from three to four. The resolution improvement obtained with the regular pixel reassignment (APR-ISM) is a less dramatic factor of *∼* 1.5. The numerical values are in good agreement with those obtained in [24, 25].

**Fig. 4.**
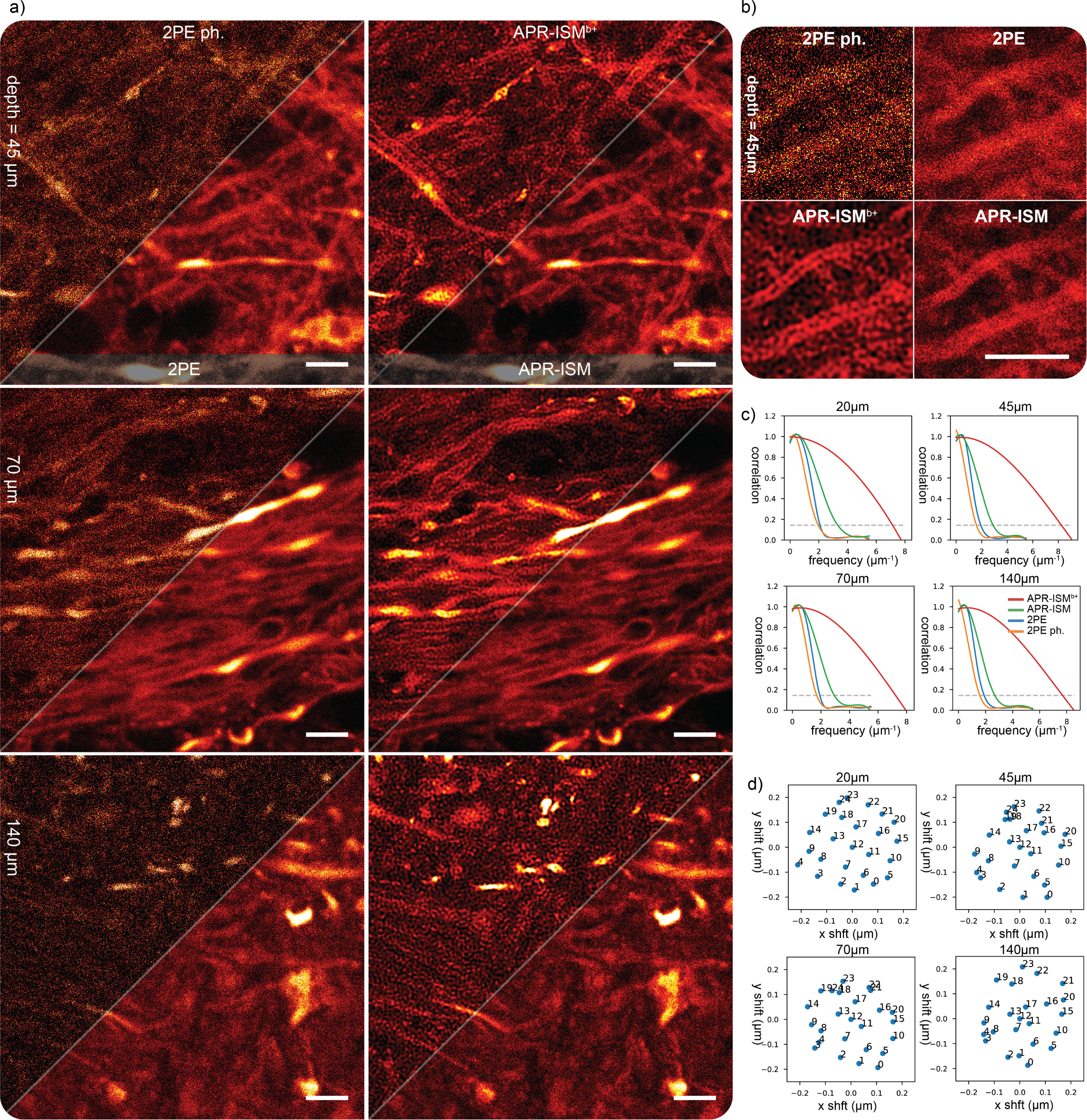
2PE-ISM imaging of a mouse brain. In a) images are shown at different penetration depths (45 *µm* – 140m *µm*), for two-photon (2PE), two-photon with pinhole (2PE ph.), adaptive pixel reassignment (APR-ISM) and adaptive pixel reassignment with blind Wiener filtering (APR-ISM^*b*+^). In b) a magnified section is shown of the results at 45 *µm* depth. In c) FRC measures for the four types of images are compared at different imaging depths. In c) 2D scatter plots of the registration results (ISM shifts) are shown. The result at 20 *µm* depth that is referred to in c, d) is shown in (Fig. S. 6). Scale bars: 4 *µm*

Considering the ISM shift vectors, our adaptive blind pixel reassignment appears to be doing much more than simply introducing a tiny virtual pinhole (Fig. 4 c). As discussed in [15], the light distribution on the array detector is related to the PSF of the microscope, and thus changes in the PSF due to different sorts of aberrations will become visible as changes in the image shifts. Typically, such aberrations are corrected during imaging with adaptive optics [25]; notably, our adaptive ISM image reconstruction appears to correct some aberrations at the post-processing stage.

## Conclusion

In this paper, we introduced a 2PE-ISM system based on a SPAD array detector that, compared to current state-of-the art, is wonderfully simple: one can convert a regular de-scanned 2PE microscope into a 2PE image-scanning microscope. In addition, thanks to the novel image reconstruction method, our 2PE-ISM microscope is very similar to use as a regular 2PE system, because no calibration measurements or other additional steps are required to achieve the improved resolution and SNR. While in this paper we focused on 2PE fluorescence imaging, the proposed architecture should also work in other non-linear (label-free) microscopy methods, such as second harmonic generation (SHG) imaging [24] and scanning laser ophthalmoscopy [40]. Furthermore, the single-photon ability of the SPAD array can be used to correlate 2PE-ISM with fluorescence lifetime imaging [15] or to combine 2PE with quantum ISM [41] to further improve the spatial resolution. We demonstrated how our adaptive blind ISM image reconstruction is able to generate nearly constant quality, high resolution images, with different samples, and at various penetration depths. In addition to the accurate image alignment, the Wiener filtering (deconvolution) appears to be an important step, as otherwise high spatial frequencies remain too weak to be observed. In this work a very simple Wiener filtering approach was used; implementing more robust (multi-image) reconstruction methods that leverage the 25 independent observations may in the future lead to further gains in image contrast and effective resolution.

Our adaptive approach can be also considered as the starting point of a new class of adaptive-optics schemes in which the SPAD array, similar to a wave-front sensor, provides the figure of merit to drive adaptive elements, such as spatial-light modulators or digital-mirror devices, able to compensate for optical aberrations. Indeed, the SPAD array provides access to information on the imaging conditions otherwise discarded by the inevitable spatial averaging effects of single-point detector.

In addition to the adaptive blind ISM image reconstruction method, a simple algorithm was proposed that allows generating a super-resolution ISM result image pixel-by-pixel for *live*-visualisation. This is an important step towards making ISM simple.

Principal limitation of our 2PE-ISM implementation is the de-scanned architecture, which does not guarantee that the scattered fluorescence photons are collected. Since the ratio between non-scattered/ballistic fluorescence photons and scattered fluorescence photons reduces for increasing depth, there will be a sample-dependent depth at which de-scanned 2PE imaging provides an effective resolution higher than our 2PE-ISM implementation. However, up to this limit non-descanned 2PE implementations can not perform better than our 2PE-ISM by increasing the intensity laser or increasing the pixel-dwell time, since the optical resolution in 2PE-ISM is always better of a factor *∼* 1.8 than regular 2PE microscopy, thus better than the de-scanning architecture.

The final thoughts, as well as several examples in this paper, help to underline the importance of computation in modern microscopy. In wide-field microscopy, similar approaches have been in regular use already for a while – e.g., axial sectioning based on deconvolution, SIM, localization based super resolution – but confocal and 2PE microscopy have remained largely analog and microscope users often have to content with the raw data. We think that ISM with its massive information content, such as our 2PE-ISM, can put an end to that.

## Methods

### The Custom Microscope Setup

Our 2PE-ISM system was implemented as a modification to a confocal microscope. As shown in Figure 2, the femtosecond two-photon excitation laser (Chameleon Ti:Sapphire, Coherent) was coupled on the confocal microscope’s common optical path via dichroic mirror DM*1 (720 *nm* SP, Semrock, USA). The fluorescence signal was directed to the SPAD array with a second dichroic mirror DM*2 (720nm LP, Semrock, USA). Both DM*1 and DM*2 are removable/exchangeable, which allows using the same microscope for single-photon ISM and regular confocal (spectroscopy) imaging. The lens pairs SL-L5 and L3-L4 conjugate the SPAD array with the object plane and adjust the magnification (*∼* 500*×*), to give the SPAD array *∼* 1.5 Airy unit field-of-view. A (512*/*40 *nm*, Semrock, USA) emission filter was installed in front of the SPAD array to block ambient light. The point-scanning was implemented using a galvanometric mirror XY scanner (6215H, Cambridge Technology, USA), coupled with *f* = 50 *mm* Leica scan lens and a *f* = 250 *mm* Leica tube lens. Axial scanning, as well as sample focusing, was implemented via a piezo stage (NanoMax MAX302, Thorlabs, USA). A single Leica Plan-Apochromat 100x/1.4-0.7 Oil CS (Leica Microsystems, Germany) objective was used in all experiments. The image acquisition was performed with our *Carma* microscope control software [42, 43], which takes care of the real-time hardware control tasks (scanning, laser control etc.) as well as fluorescence signal recording from the 25 SPAD array pixels; the software also has a PC user interface for controlling the various functions of the microscope system and to preview and process the imaging results.

### Point-spread functions simulation

The point-spread function (PSF) *h*_*i*_ associated to the *i*-th element of the detector array is calculated following the most conventional model for confocal microscopy:

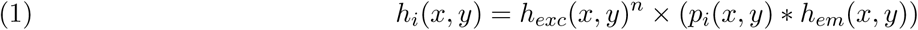

where: *h*_*exc*_ and *h*_*em*_ are the excitation and emission PSF, respectively; *p*_*i*_ is the function describing the virtual pinhole associated to the *i*-th element. In a nutshell, a binary squared mask which encodes the size of the detector element and its position; *n* is the number of photon for the excitation, i.e., *n* = 2 for 2PE and *n* = 1 for 1PE; the operator *\** denote the convolution operator. The emission and excitation PSF are calculated using a rigorous model for the calculation of the intensity distribution at the focus of an objective lens [44]. The PSF for confocal microscopy is identical to the PSF of the central element, i.e., *h*_12_. The PSF for the pinhole-less microscopy is obtained by summing all the element PSFs. The PSF for ISM is calculated implementing the pixel-reassignment method on all the PSFs *h*_*i*_.

### The ISM reconstruction method

The ISM image reconstruction is a two step process. First, all the sub-images (array pixels 1-25) need to be registered. Second, the registered sub-images need to be fused to produce a single result image.

In iterative image registration one image, called *fixed image*, is used as a reference, while a second image, called *moving image*, is translated until the details in the two images match. In our ISM registration implementation the central detector element image is always used as the *fixed image* and the 24 other images are sequentially registered with it. A rigid body spatial transformation was used, without rotation, which means that the registration entails the optimisation of two parameters. Higher level spatial transformations are supported as well in our open-source MIPLIB software library (see Acknowledgements), which may become useful for future experiments, e.g. in correcting strong aberrations. Deformable spatial transforms, in context of camera-based ISM were already discussed in [33]; the proposed approaches naturally only work if separate images are acquired for every sampling position, which typically is not the case in all-optical ISM implementations.

The ISM image registration methods in our MIPLIB library leverage the Insight Toolkit (ITK) [45], a large open-source medical image processing toolkit. The image registration pipeline is divided into several components, each of which can be selected to suit the needs of the specific task: *metrics, optimizers, transforms, interpolators* and *initializers*. The *metric* is used to measure the similarity of the *moving* and *fixed* images; the aim of a registration task is to maximize its value. The *optimizer* adjusts the *transform* parameters, until the similarity *metric* reaches its maximum value. The *interpolator* is used to calculate pixel values at non-grid positions during transformation. The *initializer* calculates the initial transformation for rough alignment of the *moving image* with the *fixed image*. The components of our ISM image registration implementation are:

> **Metric:** Normalised Cross Correlation that is calculated from 0.01-1% subset of the pixel values; a higher percentage is required for low quality or extremely sparse images. Different similarity metrics are also supported in the MIPLIB software.
>
> **Optimizer:** Regular Step Gradient Optimizer, which essentially at each iteration takes a step along the direction of the metric derivative
>
> **Initializer:** not needed (images are already sufficiently overlapped); both automatic and manual offset initialization is supported in MIPLIB software.
>
> **Transform:** 2D rigid translation transform. Scalable and deformable spatial transformations are supported as well, but not used here.
>
> **Interpolator:** linear interpolation

All registration and image transformation tasks in ITK/MIPLIB are performed in physical units (*µm*), which means that the images can be re-sampled to a different pixel size without changing the registration result. In ISM this is particularly useful, if the original data is sampled too sparsely to support the two-fold resolution improvement. One can also shrink the images before registration to increase speed and optimise memory consumption. The registration is also always sub-pixel: for the results shown in this work, the minimum *optimizer* step size was set to 0.5 nm.

After all the 25 images have been registered, they can be added together to for the regular APR-ISM pixel reassignment result. In order to take full advantage of ISM however, one has to still remove the blurring effect of the PSF, which in this case is that of the ISM reassignment result, as explained in context of SIM image reconstruction in [39]. To this end, we applied our FRC based blind Wiener filter on the reassignment result. The filter first estimates the PSF directly from the image with an FRC measurement, after which a classical Wiener filter is applied; the FRC cut-off value is used as a full-width-half-maximum (FWHM) value for a Gaussian PSF. Please refer to [38] for a detailed description.

The ISM image reconstruction workflow described above is mainly intended for the post-processing stage. The pixel reassignment can, however, also be performed in real-time, pixel-by-pixel, just as in a regular confocal or two-photon microscope (Note S 1).

### Resolution measurement

The resolution measurements shown in this work, were calculated with our Fourier-ring-correlation (FRC) assay [46]. When two “identical” images (i.e, image of the very same structures, but with different noise realisation) were not available, we used a one-image version of the FRC assay. The principle is the same as with the blind image deconvolution: first an image is split into two sub-images, after which the FRC is taken with the two sub-images as inputs. The one-image FRC method is described in detail in [38]

### Test Samples

We demonstrated the enhancement on spatial resolution of 2PE-ISM *via* imaging of fluorescent beads, tubulin filaments and optically cleared mouse brain.

> **Fluorescent beads:** In this study we used Yellow/Green fluorescent nanoparticles with a diameter of 100 nm (FluoSpheres, ThermoFisher Scientific, USA).
>
> **Tubulin filaments in fixed cells:** Human HeLa cells were fixed with ice-cold methanol for 20 ° min at 20 ° C and then washed three times for 15 min in PBS. After 1 h at room temperature, the cells were treated in a solution of 3% bovine serum albumin (BSA) and 0.1% Triton in PBS (blocking buffer). The cells were then incubated with monoclonal mouse anti-*α*-tubulin antiserum (Sigma-Aldrich) diluted in blocking buffer (1:800) for 1 h at room temperature. The *α*-tubulin antibody was revealed by Abberior STAR488 goat anti-mouse (Abberior) for the custom microscope. The cells were rinsed three times in PBS for 5 min.
>
> **Optically cleared brain of Thy1-eYFP-H transgenic mouse:** The CLARITY method was used to clear the mouse brain [47]. In short, after perfusion, mouse brains were post-fixed in 4% PFA overnight at 4 °C and then immersed in 2% hydrogel (2% acrylamide, 0.125% Bis, 4% PFA, 0.025% VA-044 initiator (w/v), in PBS) for 3 days at 4 °C. Samples were degassed and polymerized for 3.5 hours at 37 °C. The samples were removed from hydrogel and washed with 8% SDS for 1 day at 37 °C. The samples were transferred to fresh 8% SDS for 21 days at 37 °C for de-lipidation. Then the samples were washed with 0.2% PBST for 3 days at 37 °C. Brains were incubated in RapiClear CS (Cat#RCCS002, SunJin Lab) for 2-3 days at room-temperature for the optical clearing.

## Acknowledgements

We thank Ryu Nakamura (Nikon Corporation, Japan) for support on brain imaging. We thank Ryu Nakamura (Nikon Corporation, Japan), Dr. Ryosuke Kawakami, Dr. Kohei Otomo, and Prof. Tomomi Nemoto (Research Institute for Electronic Science, Hokkaido University, Japan) for advise in whole brain imaging. We thank Dr. Michele Oneto (Istituto Italiano di Tecnologia, Italy) for support in cell preparation, Dr. Simonluca Piazza (Istituto Italiano di Tecnologia, Italy) for useful discussions.

## Author contributions

S.K., E.S., and G.V. conceived the idea. S.K. and G.V. supervised the project, with support from C.J.R.S., A.D., and A.T.. E.S., S.K., E.T., G.T., P.B., and G.V. designed and implemented the custom 2PE-ISM system. S.K., G.T., and M.C., developed the image processing and analysis software. G.T. performed the simulations. E.S. performed the experiments. S.K. and G.V. analysed the data with support from all authors. M.C., G.T. and G.V. designed and developed the controlling architecture. M.C., M.B., F.V., A.T., and G.V., designed the SPAD array module. M.B., F.V., and A.T. realised and characterised the SPAD array module. S.K., E.S., and G.V. wrote the manuscript with input from all authors.

## Code Availability

All the image analysis and processing methods used in this work are available as a part of the MIPLIB software library at (https://github.com/sakoho81/miplib)

## Data availability

All the data that supports the findings of this study are available from the corresponding author upon request.

## Supporting Information

**Fig. S 1.**
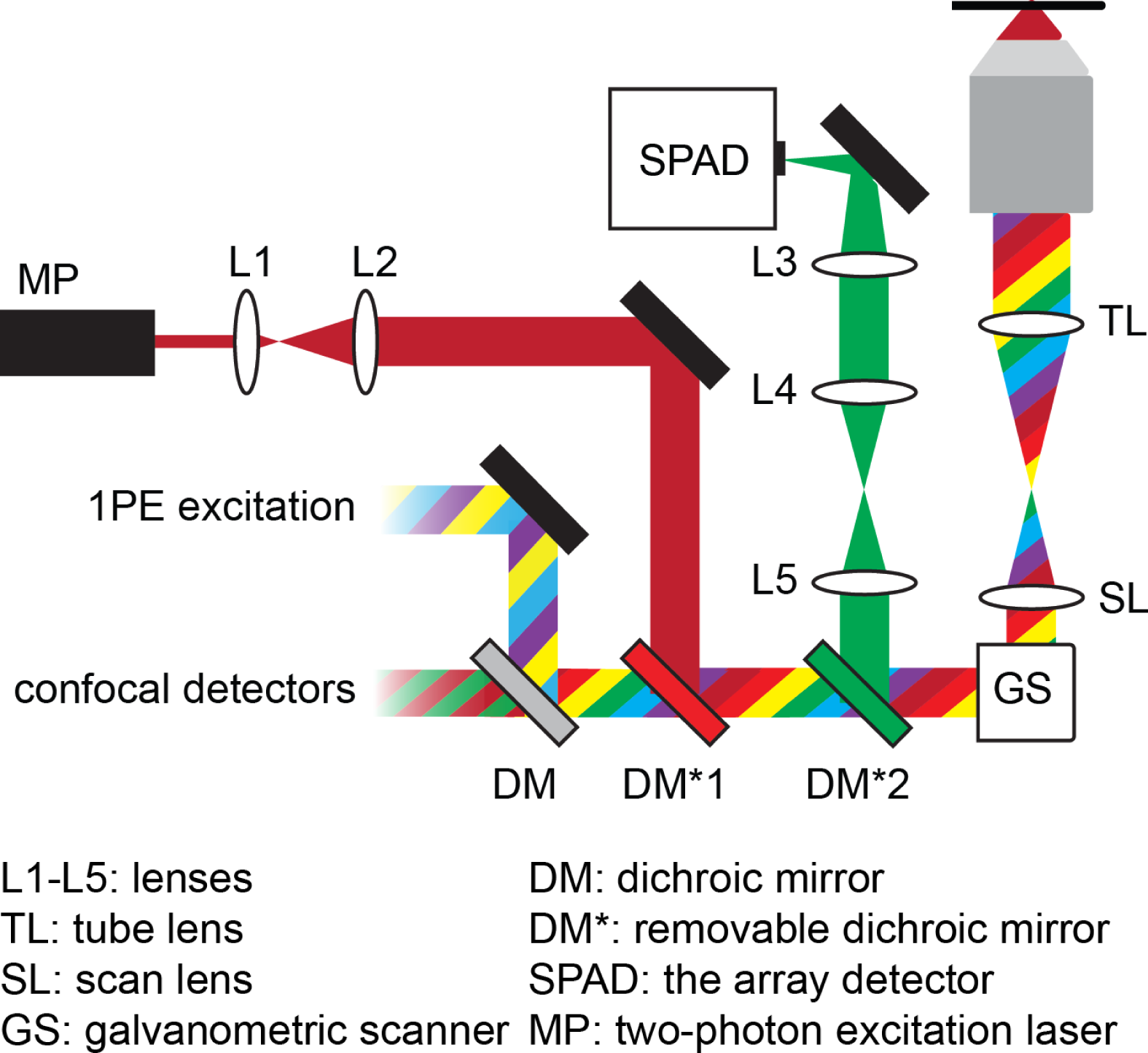
Schematic of the 2PE-ISM setup. A schematic of the 2PE-ISM system is shown. The ISM part was built as an extension to a regular confocal microscope with several laser lines and detectors. Only three additional lenses (L3-L5) and a dichroic mirror (DM2) are needed to convert a confocal/2PE system into a ISM super-resolution microscope. The dichroic mirrors DM1-2 can be reconfigured to enable ISM imaging with the visible excitation lines.

**Fig. S 2.**
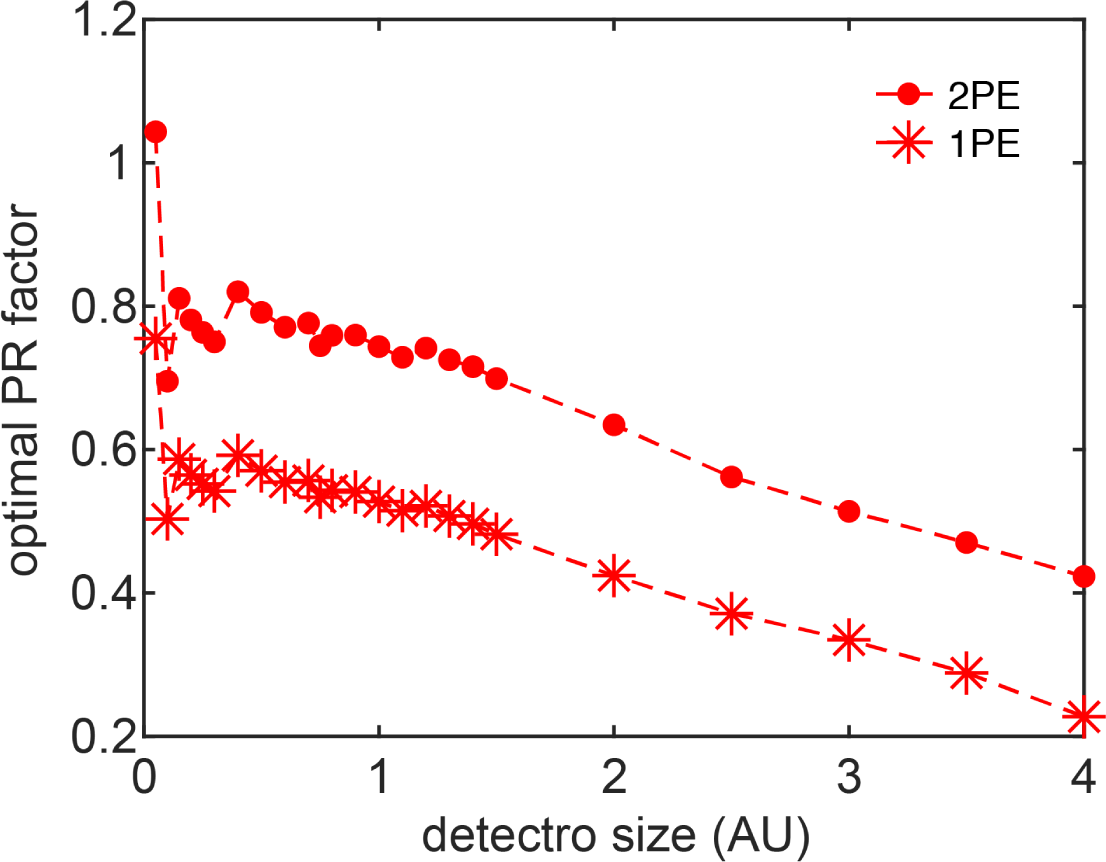
Optimal PR factor. Calculated PR factor as a function of the detector size for the 2PE and 1PE cases. The PR factor are calculated by using the shift-vector derived from our PSF simulation. Here, the shift-vectors are derived assuming that the optimal ISM reconstruction is obtained by co-aligning all the element PSFs to their maximum intensity before their summation. After all shift-vectors are obtained, the average module 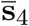 for the first-neighboured elements is calculated 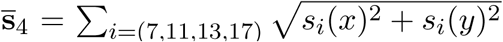. This average module multiplied by the system magnification and divided by the pixel-pitch of the detector gives the optimal PR factor.

**Fig. S 3.**
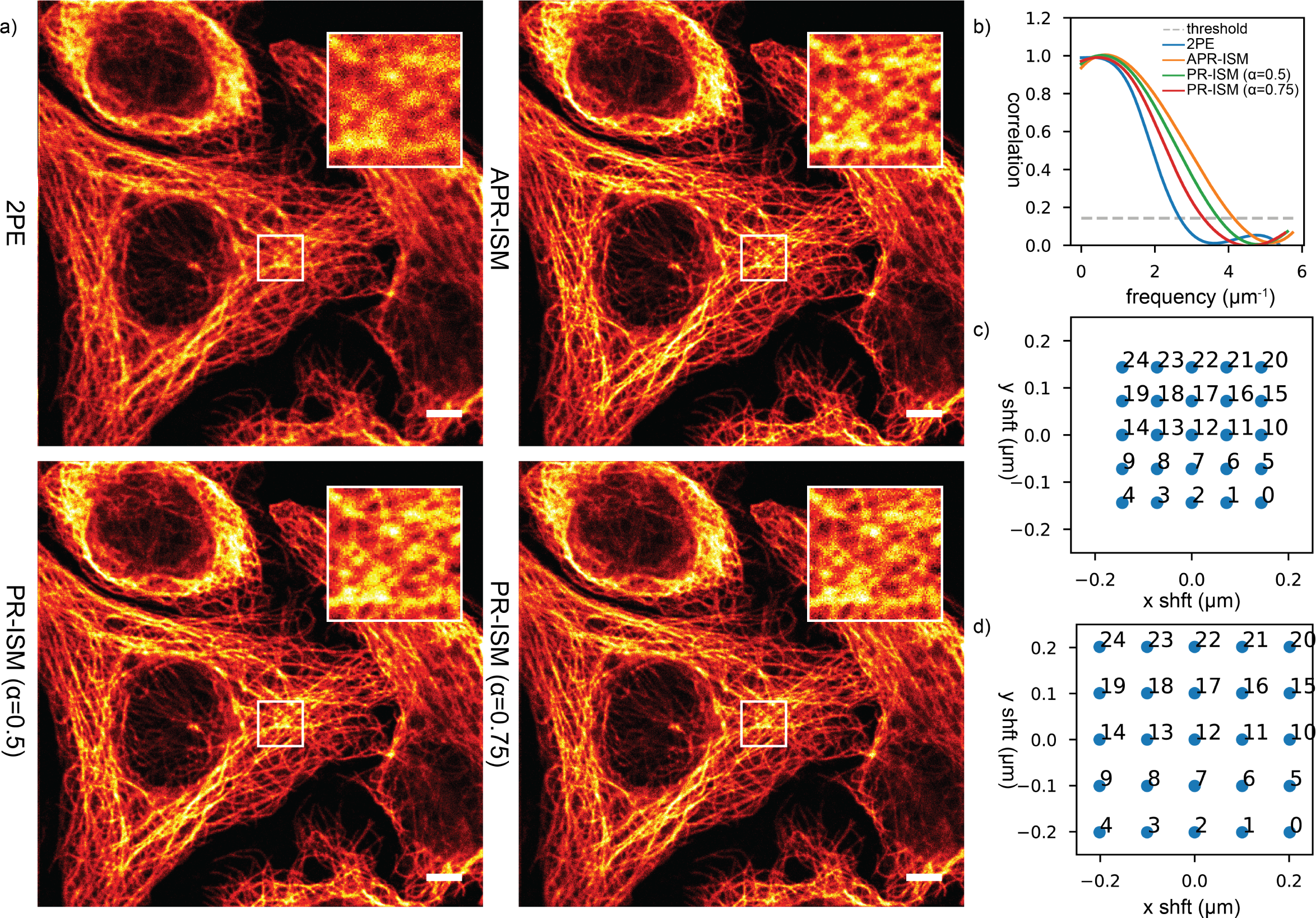
Comparing ISM image reconstructions with adaptive and theoretical shift-vectors. ISM reconstructions are shown for the HeLa cell image with adaptive and theoretical shift-vectors. Two value for the pixel-reassignment (PR) factor are tested, *α*=0.5 and *α*=0.75, respectively the value used in many all-optical ISM implementations and the value obtained from our simulation. The adaptive ISM reconstruction produces clearly superior results. Scale bar 4 *µm*.

**Fig. S 4.**
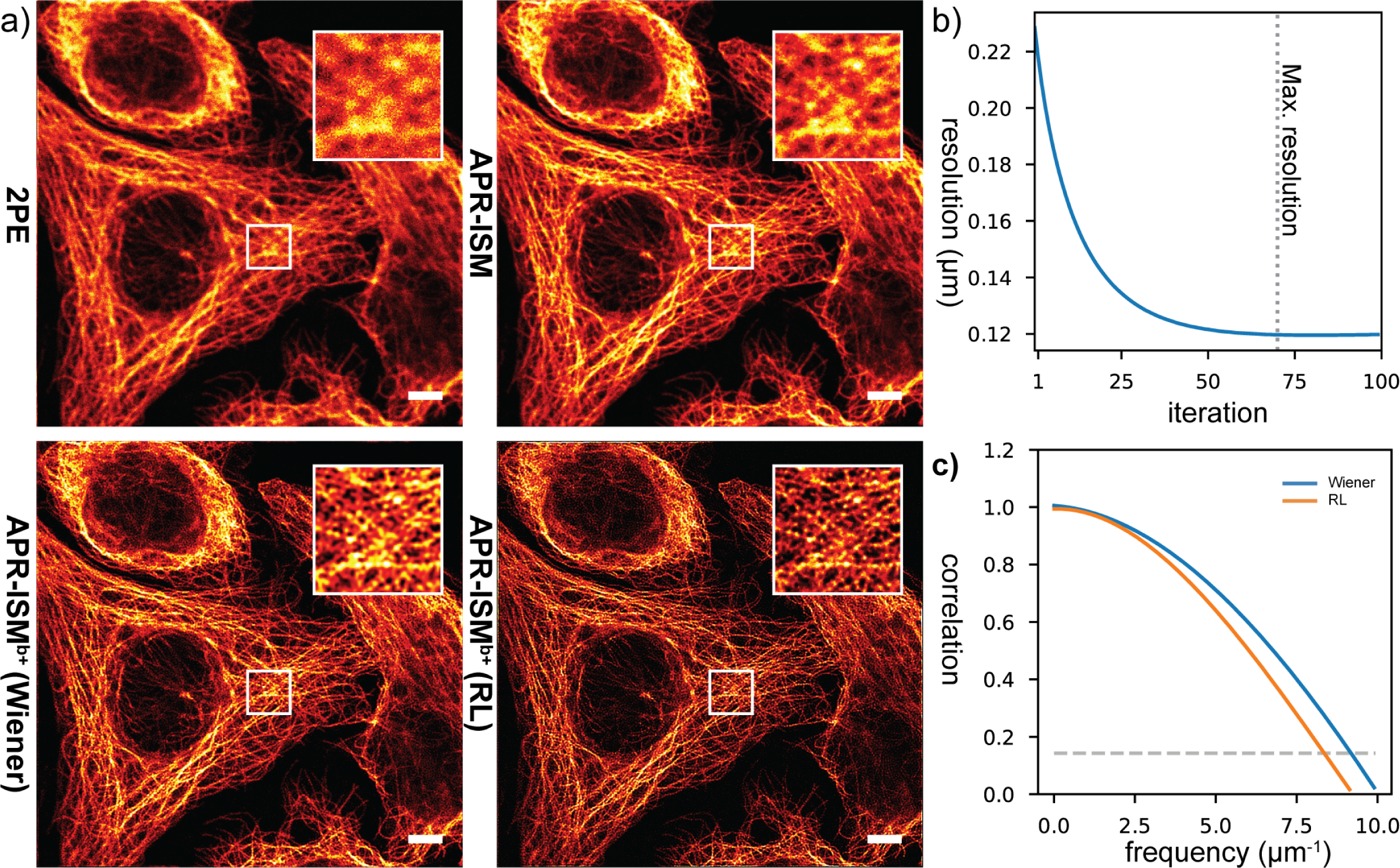
Comparing blind Wiener and Richardson Lucy deconvolution. In a) the performance of the blind Wiener filtering is compared against iterative Richardson-Lucy (RL) deconvolution. The RL algorithm produces sharper looking results with strongly improved contrast, but quantitatively, as shown by FRC measurements in c), the resolution in the two images is the same. It may thus be beneficial to use RL or other iterative algorithm to produce the crispiest looking results, but as shown in b) it takes about 50 iterations for the RL algorithm to reach the same resolution scale the the Wiener filter is able to produce with a single division. Thus, when speed is an issue, Wiener filter provides a much superior solution (of course RL can be significantly accelerated with GPU if necessary). Scale bar 4 *µm*.

**Fig. S 5.**
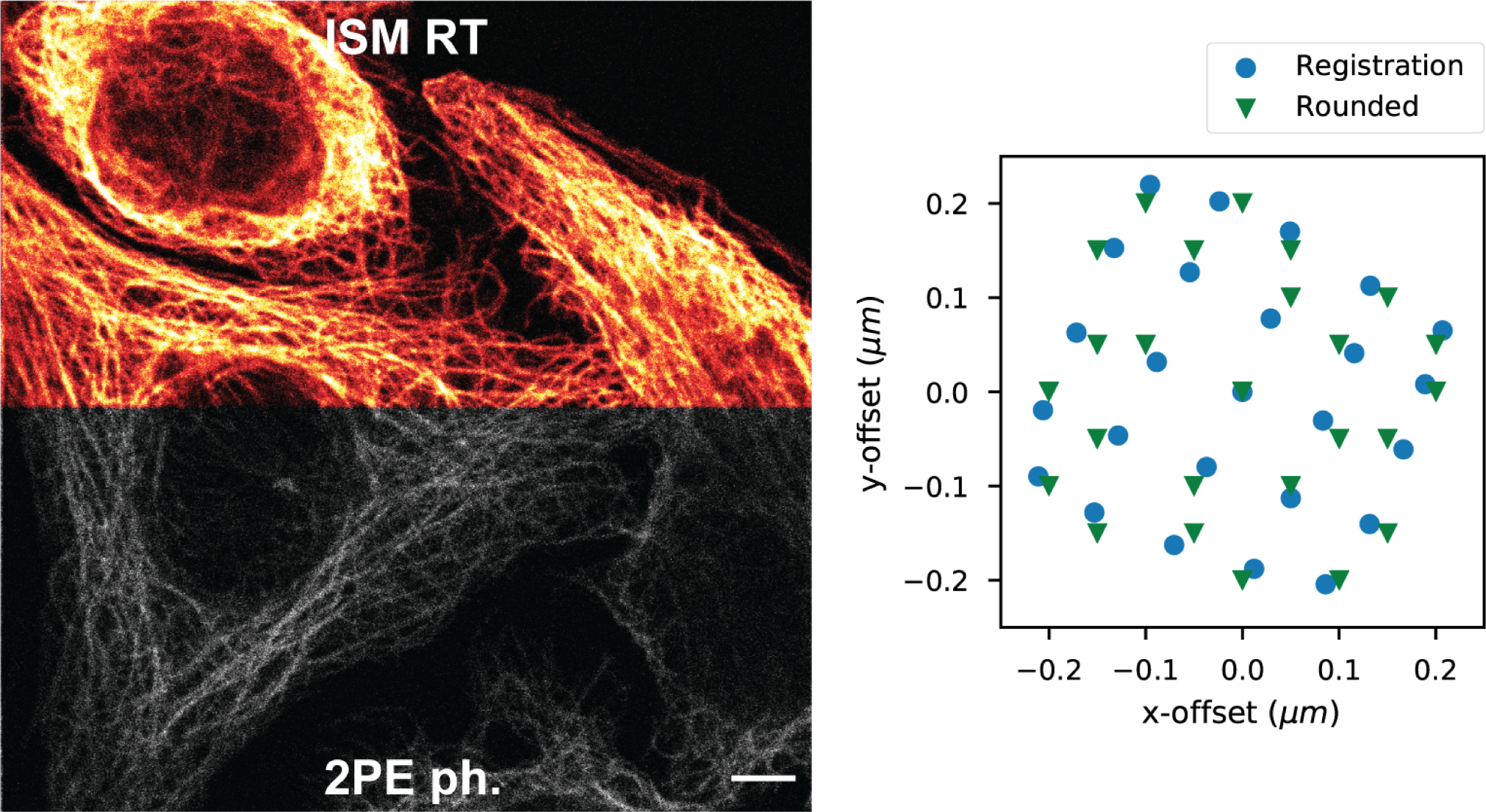
Real-time pixel-reassignment in action. Real-time pixel reassignment is shown to dramatically boost the SNR in the HeLa cell image, when compared to 2PE image with a pinhole. This makes it possible to achieve the resolution gain (and optical sectioning capability) in practice that the small pinhole can in theory provide. In the scatter plot ISM shifts discretized to multiples of the pixel size (Note S 1) are compared to the actual registration results. The discretized shifts are used by the real-time pixel reassignment algorithm, because no re-sampling is performed, but simple array indexing is used instead. While the rounding does produce small errors, the observed image quality still remains high. Scale bar 4 *µm*.

**Fig. S 6.**
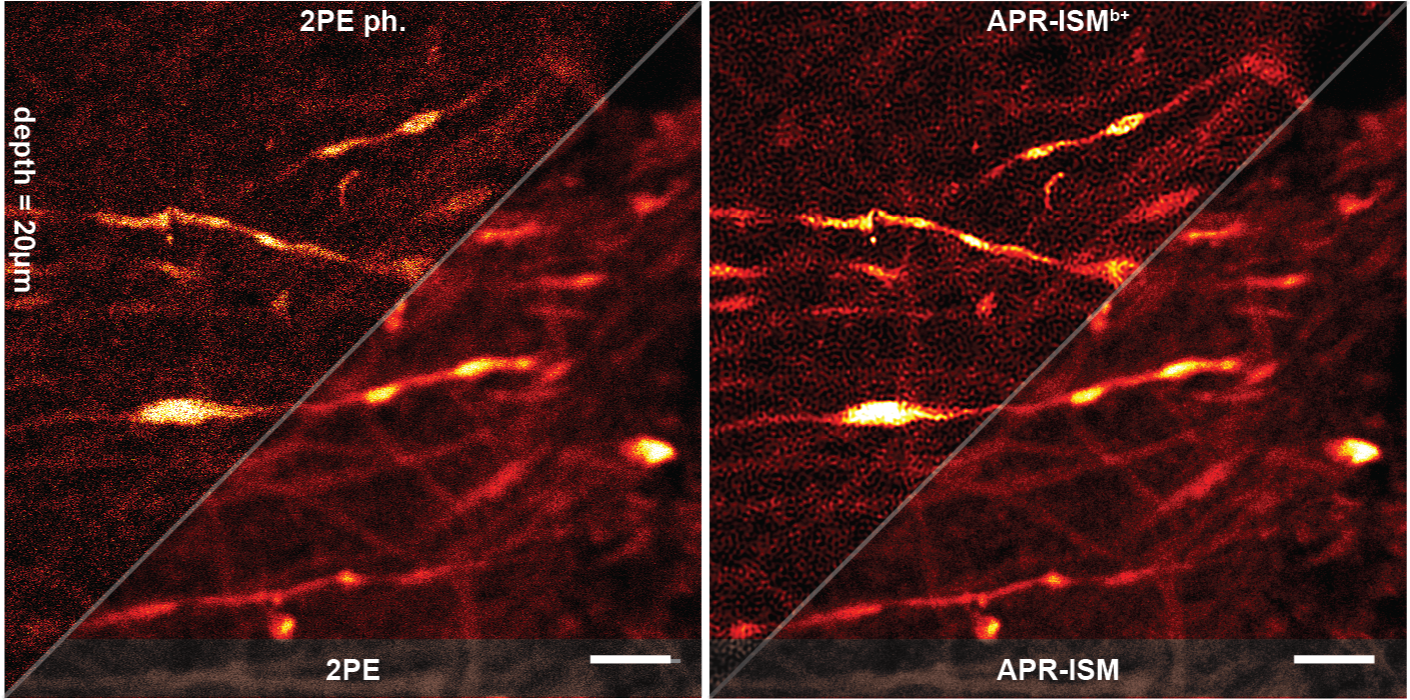
ISM results at 20 *µm* depth in mouse brain. ISM reassignment results at 20 *µm* depth in the brain sample are shown. Scale bars 4 *µm*.

## Note S 1. Real-Time ISM Pixel Reassignment Algorithm

Our real-time pixel reassignment algorithm is shown in (Algorithm S 1). Image of size (*imageWidth, imageHeight*) is formed one sampling position at a time, typically by laser-scanning. At each sampling position (*column, row*) the photon counts from each pixel of the array detector are added to the correct position in the *resultImage*, by applying the image shifts (*shiftsY, shiftsX*). The image shifts can be based on image registration results of the previous frames, or alternatively theoretical values can be used. In order to perform the reassignment in real-time, the (*shiftsY, shiftsX*) need to be expressed as pixels in stead of physical distances, which means that they have to be rounded to multiples of the pixel size. This can create a small error in the shifts as shown in (Fig. S5), but in most cases this should not be an issue. It is possible to scale the result image to a smaller pixel size than the sampling grid, if necessary, to account for the higher resolution and to decrease the shift error. In (Algorithm S 1) this would simply involve creating a larger result image, and scaling the shifts by the ratio of the sampling grid and the result image pixel sizes.

### Algorithm S 1 Simple pseudocode for our real-time pixel reassignment algorithm. Words in italics denote variable names

*resultImage ←* zeros((*imageHeight, imageWidth*))

**for** *column* in **range**(*imageWidth*) **do**

**for** *row* in **range**(*imageHeight*) **do**

*arrayData ←* **GetPhotonCounts**()

**for** *detector* in **range**(*nDetectors*) **do**

*pixel ← arrayData*[*detector*]

*xIdx ←* **int**(*column-shiftsX*[*detector*])

*yIdx ←* **int**(*row-shiftsY* [*detector*])

**if** *xIdx ≥* 0 **and** *xIdx < imageWidth* **and** *yIdx ≥* 0 **and** *yIdx < imageHeight* **then**

*resultImage*[*yIdx, xIdx*] += *pixel*

**end if**

**end for**

**end for**

**end for**

